# *Anopheles barbirostris* in Indonesia: A more complex metapopulation than expected

**DOI:** 10.1101/2025.03.12.642759

**Authors:** Tri Baskoro Tunggul Satoto, Triwibowo Ambar Garjito, Shinta, Soleman Landi, Roger Frutos, Sylvie Manguin

**Affiliations:** Department of Parasitology, Faculty of Medicine, Public Health and Nursing, Gadjah Mada University, Yogyakarta, Indonesia; Research Center for Public Health and Nutrition, National Research and Innovation Agency Indonesia; Public health faculty, Nusa Cendana University, East Nusa Tenggara; Cirad, UMR 17, Intertryp, Montpellier, France; Faculty of Medicine-Ramathibodi Hospital, Mahidol University, Bangkok, Thailand; Department of Health, Faculty of Vocational Studies, Universitas Airlangga, Surabaya, Indonesia; HSM, University of Montpellier, CNRS, IRD, Montpellier, France

**Keywords:** *Anopheles barbirostris* complex, *Anopheles wejchoochotei*, molecular markers, ITS2, COI, genetic diversity, Indonesia

## Abstract

*Anopheles barbirostris*, a member of the Barbirostris Subgroup in the *Anopheles* genus, comprises a complex of species in South and Southeast Asia. This *An. barbirostris* complex includes eight species such as *An. barbirostris sensu stricto* (s.s.), *An. campestris, An. dissidens, An. donaldi, An. saeungae, An. sarpangensis, An. vanderwulpi*, and *An. wejchoochotei*. This study employed molecular markers, including ITS2 and COI genes, to investigate the phylogenetic relationships within *An. barbirostris sensu lato* (s.l.) populations from various locations in Indonesia. The analysis reveals the presence of nine distinct populations within this complex, including the first report of *An. wejchoochotei* in North Sulawesi and an unique Barbirostris population in Magelang, Central Java. This makes it a more complex metapopulation than previously thought. These findings provide critical insights into the diversity of malaria and lymphatic filariasis vectors in Indonesia. Understanding the complex structure of the *An. barbirostris* populations and their genetic diversity will be useful for effective vector control and disease elimination strategies.

## Introduction

*Anopheles barbirostris* is currently identified as a species within the Barbirostris Subgroup of the Subgenus *Anopheles* [1,2]. This subgroup is composed of 11 species, including six that are differentiated by distinct morphological characters, such as *An. barbirostris, An. campestris, An. donaldi, An. franciscoi, An. hodgkini*, and *An. pollicaris* [3,4]. *Anopheles barbirostris sensu lato* (s.l.) has been regarded as a species complex since 2001 according to cytogenetics and molecular characteristics [2,5,6]. This *An. barbirostris* complex includes eight species such as *An. barbirostris, An. campestris, An. dissidens, An. donaldi, An. saeungae, An. sarpangensis, An. vanderwulpi*, and *An. wejchoochotei* [2,4,6,7]. This species complex is broadly distributed in South and Southeast Asia, such as Indonesia, Thailand, Malaysia, Timor Leste, Vietnam, Myanmar, Cambodia, Sri Lanka, Nepal, Bhutan, Bangladesh, and China [3,4,8–13]. In Indonesia, *An. barbirostris* s.l. has been reported in Sumatra, Java, Kalimantan, Sulawesi, Bali, Lesser Sunda Islands, and North Maluku (Buru Island). However, this species complex has never been reported from the Maluku and Papua regions [9,14].

Prior to the implementation of molecular analyses, *An. barbirostris* s.l. was regarded as comprising 4 distinct cytological forms [15]. The first evidence of *An. barbirostris* as a species complex was formally reported as W, X, and Z forms based on mtDNA Cytochrome Oxidase I gene (COI) [5]. Subsequent molecular phylogenetic analyses of *An. barbirostris* s.l. based on mtDNA COI and rDNA ITS2 have revealed the presence of several sympatric clades in Sumatra and Java [3]. More recently, based on integrated morphological and molecular (COI and ITS2) characterization, four species within the Barbirostris complex have been reported in Indonesia: *An. barbirostris* in Java and Kalimantan, *An. vanderwulpi* in Java and Sumatra, *An. saeungae* in Sumatra, and an unknown form of *An. barbirostris* in Sulawesi [3,8,16– 18].

*Anopheles barbirostris* s.l. has been considered an important malaria and lymphatic filariasis vector due to its anthropophilic behavior in Lesser Sunda Islands and Sulawesi [19–21]. *Anopheles barbirostris* s.l was first reported as a malaria vector in 1939 in South Sulawesi [22]. The role of this taxon as an important malaria vector was also reported in several locations in the Lesser Sunda Islands (Lombok, Flores, and Adonara Islands), Northern Sulawesi (Meras and Tomohon), and Central Sulawesi [23]. Both *Plasmodium falciparum* and *Plasmodium vivax* infections were detected in all of these areas [19,21,24,25]. *Anopheles barbirostris* s.l. has also been confirmed as a Malayan filaria vector in Sulawesi and as a Timor filaria vector in East Nusa Tenggara [26,27]. However, while present in Sumatra, Java, Kalimantan, West Nusa Tenggara, and Maluku, *An. barbirostris* s.l. has not been confirmed as malaria and/or lymphatic filariasis vector in these areas due to its mainly zoophilic behavior [9]. Therefore, it is of upmost importance to identify the species of the Barbirostris complex to evaluate their behavior and their involvement in parasite transmission.

In this study, we used the internal transcribed spacer 2 (ITS2), a ribosomal DNA (rDNA) region, and the cytochrome oxidase I (COI) mitochondrial gene to characterize the phylogenetic relationships of the *An. barbirostris* s.l. samples collected from different locations in Indonesia. This information is a first but important step towards the understanding of local malaria transmission dynamics and the design of appropriate vector control strategies within the national malaria elimination program.

## Material and Methods

### Mosquito field collections and species identification

Adult mosquitoes were collected from the field using human-landing collection (HLC) and cattle-baited trap methods in the provinces of North Sulawesi, Central Sulawesi, South Sulawesi, Central Java, East Java, and East Nusa Tenggara (Table 1, Fig. 1). All specimens were initially identified as *An. barbirostris* s.l. based on standard morphological adult mosquito identification keys for Indonesia [28,29]. These mosquitoes were labeled according to locality and date, and kept individually in 1.5 ml tubes under dry conditions using silica gel until molecular analysis was performed.

**Table 1.** Sampling localities and specimens of the *Anopheles barbirostris* complex.

| Sample code | Province | Location | Malaria vector | GenBank ID (ITS2) | GenBank ID (COI) |
| --- | --- | --- | --- | --- | --- |
| 1-2 | South Sulawesi | Pangkajene Kepulauan | Yes | X | X |
| 2-2 | South Sulawesi | Pangkajene Kepulauan | Yes | X | X |
| 3-2 | Central Sulawesi | Donggala | Yes | X | X |
| 4-2 | Central Sulawesi | Donggala | Yes | X | X |
| 5-2 | Central Sulawesi | Donggala | Yes | X | X |
| 6-2 | Central Java | Semarang-Salaman- Jambu | No | X | X |
| 7-2 | East Java | Malang | No | X | X |
| 8-2 | East Java | Malang | No | X | X |
| 9-2 | East Nusa Tenggara | Flores | Yes | X | X |
| 10-2 | Central Java | Magelang | No | X | X |
| 11-2 | North Sulawesi | Manado | Yes | X | X |
| JAVA-J7 | Central Java | Semarang | No |  | X |
| JAVAS8 | Central Java | Magelang | No |  | X |
| JAVAJ10 | Central Java | Semarang | No |  | X |
| FLORES-S1 | East Nusa Tenggara | Flores | Yes |  | X |
| FLORES-S3 | East Nusa Tenggara | Flores | Yes |  | X |
| FLORES-R8 | East Nusa Tenggara | Flores | Yes |  | X |

**Figure 1.**
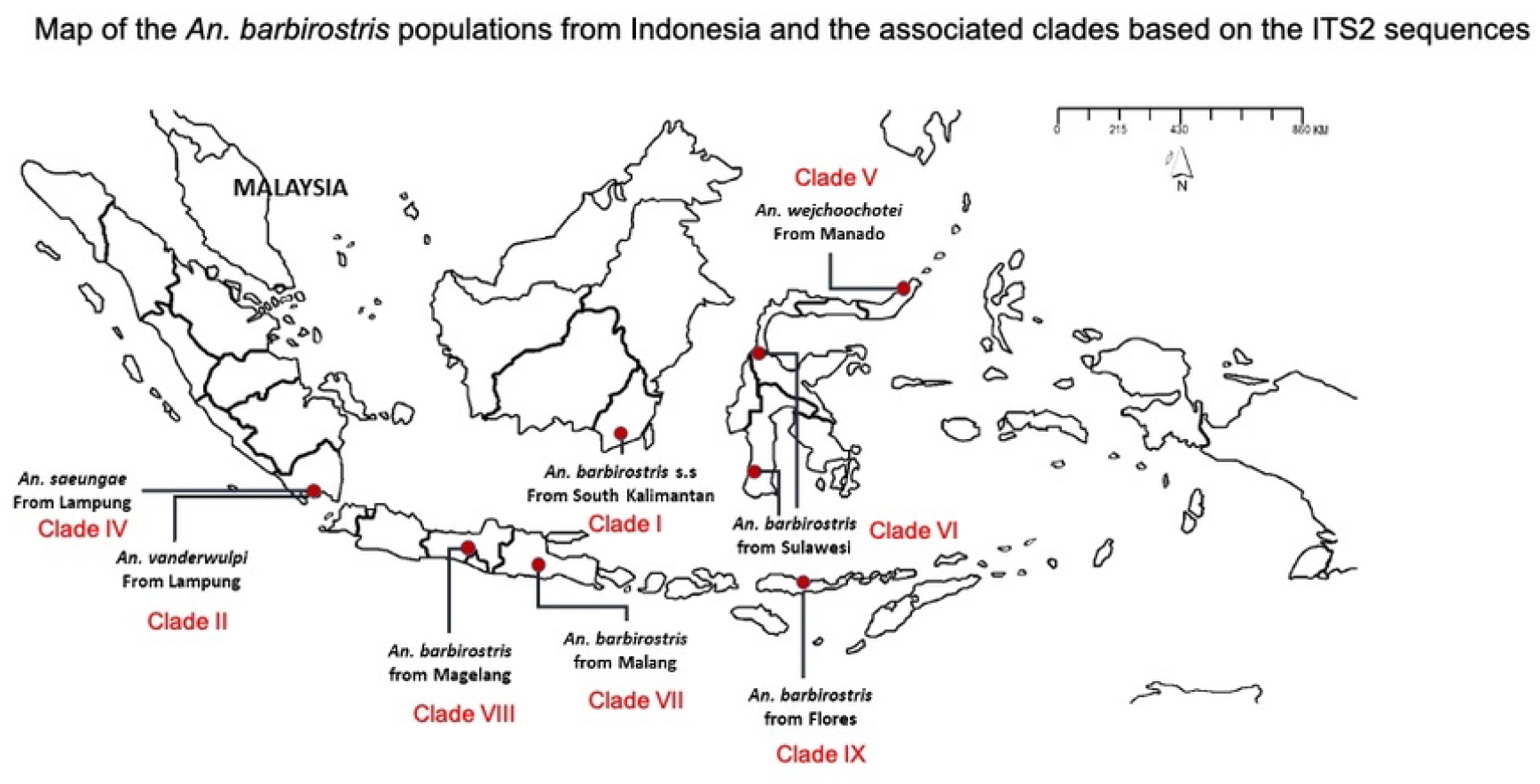
Map of the *An. barbirostris* s.l. associated to 8 clades (I-II, and IV-IX) based on the ITS2 sequences, and locations of the sampling sites (red spots) in Indonesia.

### rDNA-ITS2 and mtDNA-COI gene amplification and sequencing analysis

The genomic DNA was extracted from individual mosquitoes using a DNeasy^®^ Blood & Tissue Kit (Qiagen, Hilden, Germany) based on the manufacturer’s protocol. The amplification of rDNA-ITS2 was performed with universal primers ITS2-F (5’-GGT GGT GAC TTT CAA GTTC-3’) and ITS2-R (5’-TAT GCT TAA ATT TAG TTT GTA G -3’). PCR reactions were performed using GoTaq^®^ Green Master Mix (Promega Madison, WI, USA). PCR thermocycling conditions for ITS2 were as follows: 95°C for 5 min; followed by 26 cycles of denaturation at 94°C for 1 min, annealing at 55°C for 1 min and elongation at 72°C for 1 min; and followed by a final extension step at 72°C for 7 min. The mtDNA-COI was amplified using primer CI-N-2087 (5’-AAT TTC GGT CAG TTA ATA ATA TAG-3’) and TY-J-1460 (5’-TAC AAT TTA TCG CCT AAA CTT CAG CC-3’) [30]. PCR was also carried out using GoTaq^®^ Green Master Mix (Promega Madison, WI, USA). The mtDNA-COI amplification comprised initial denaturation at 94°C for 1 min followed by five cycles of 94°C for 30 s, 45°C for 40 s, and 72°C for 1 min. This process was then followed by 35 cycles of 94°C for 30 s, 55°C for 40 s, 72 for 1 min, and a final extension at 72°C for 10 min. All PCR reactions were performed in an Applied Biosystems SimpliAmp thermal cycler. After PCR amplification, products were separated by 1.5% agarose gel electrophoresis and stained with SYBR^®^ safe DNA gel stain (Invitrogen, Carlsbad, CA, USA). The amplification products were purified using Applied Biosystems ExoSAP-IT™ (Thermo Fisher Scientific, Vilnius, Lithuania) and sequenced using the primers listed above with an Applied Biosystems Big Dye™ Terminator version 3.1 Sequencing Kit (Life Technologies Cooperation, Austin, TX, USA). Cycle sequencing products were edited using Sequencing Analysis v5.2. (Applied Biosystems), and were aligned using ClustalW 1.6 in Mega X version 10.2.2. The sequences were compared with the available sequences of *Anopheles barbirostris* in the NCBI GenBank database. The maximum likelihood method with the General Time Reversible (GTR)+1 model was constructed in Mega X version 10.2.2. *Anopheles barbumbrosus*, of the Barbirostris Group, was used as an outgroup in the phylogenetic trees. Bootstraps were tested with 1,000 replicates to assess the reliability of the trees. Variations within and between species were measured by the pairwise distance (p-distance) method in MegaX version 10.2.2. Sequences are deposited in NCBI GenBank with the following accession numbers: PP840883-PP840889, PP840890, PP840892-PP840894.

## Results

### Phylogenetic analysis of the Internal Transcribed Spacer 2 (ITS2)

The analysis of the ITS2 sequences of the *An. barbirostris* samples from this work and reference sequences available from GenBank revealed that eight populations of *An. barbirostris* could be identified in Indonesia (Fig. 2). They include *An. barbirostris* from South Kalimantan (EU812759-EU812760 - clade I), *An. vanderwulpi* from Lampung-Sumatra (EU812765-EU812768 - clade II), *An. saeungae* from Lampung-Sumatra (EU812791 – clade IV), *An. wejchoochotei* from Manado-North Sulawesi (sample 11.2 – clade V), *An. barbirostris* from South (samples 1.2, 2.2) and Central Sulawesi (samples 4.2, 5.2), and Central Java (sample 6.2) all of clade VI, *An. barbirostris* from Malang-East Java (samples 7.2-8.2 – clade VII), *An. barbirostris* from Magelang-Central Java (sample 10.2 – clade VIII), and *An. barbirostris* from Flores-East Nusa Tenggara (sample 9.2 – clade IX). The ITS2 of *An. barbirostris* samples from Malang-East Java (samples 7.2, 8.2 – clade VII) differ from *An. vanderwulpi* from Sumatra (EU812765-EU812768 – clade II) by 1.3 - 2.7%. High conservation and 100% genetic similarity were identified among samples from Pangkep, South Sulawesi (samples 1.2, 2.2), samples from Semarang-Central Java (sample 6.2), and those from Donggala-Central Sulawesi (samples 4.2, 5.2), all belonging to clade VI. The sample of *An. barbirostris* from Manado, North Sulawesi (sample 11.2 – clade V) displayed 100% ITS2 identity with *An. wejchoochotei* from Thailand (AB971307, AB971309 – clade V) and also showed 99.7% genetic similarity with *An. campestris* from Thailand (EU812808-EU812809 – clade V). The Sulawesi samples (1.2, 2.2, 4.2, 5.2 – clade VI) and Central Java sample (6.2 – clade VI) displayed 0% internal group distance and distances of 3.2-4.3 % with *An. barbirostris* from Magelang-Central Java (sample 10.2 – clade VIII), 0.1-2.4% with *An. barbirostris* from South Kalimantan (EU812759-EU812760 – clade I), 3.7-4.3% with *An. barbirostris* from Flores, East Nusa Tenggara (sample 9.2 – clade IX), and 4.5-5.6% *An. barbirostris* from Thailand (EU812761-EU812764 – clade I). Furthermore, the distances between *An. barbirostris* from Lampung-West Sumatra (EU812791 – clade IV) and *An. saeungae* from Thailand (MH796417, MH796419, MH796422, MH796424, EU812794 - clade IV) were 0.7%-4.2%. Moreover, the *An. barbirostris* specimen (sample 10.2 – clade VIII) from Magelang-Central Java was also identified as a distinct species of the *An. barbirostris* complex in Indonesia. In comparison with other members in this study, the Magelang-Central Java ITS2 sequences displayed distances of 3.5%, 1.0%, 3.1-4.3%, and 4.0-4.5% with *An. barbirostris* from South Kalimantan (EU812759-EU812760 – clade I), Flores-East Nusa Tenggara (sample 9.2 – clade IX), Sulawesi (samples 1.2, 2.2, 4.2, 5.2, 6.2 – clade VI), and from Thailand (EU812761-EU812764 – clade I) respectively. The current study also suggests the presence of an additional new distinct population, i.e. *An. barbirostris* from Flores-East Nusa Tenggara (sample 9.2 – clade IX) identified as being relatively closely related to *An. barbirostris* k2 and k3 from South Kalimantan (EU812759-EU812760), although genetically distinct by 2.4% from them (Fig. 2).

**Figure 2.**
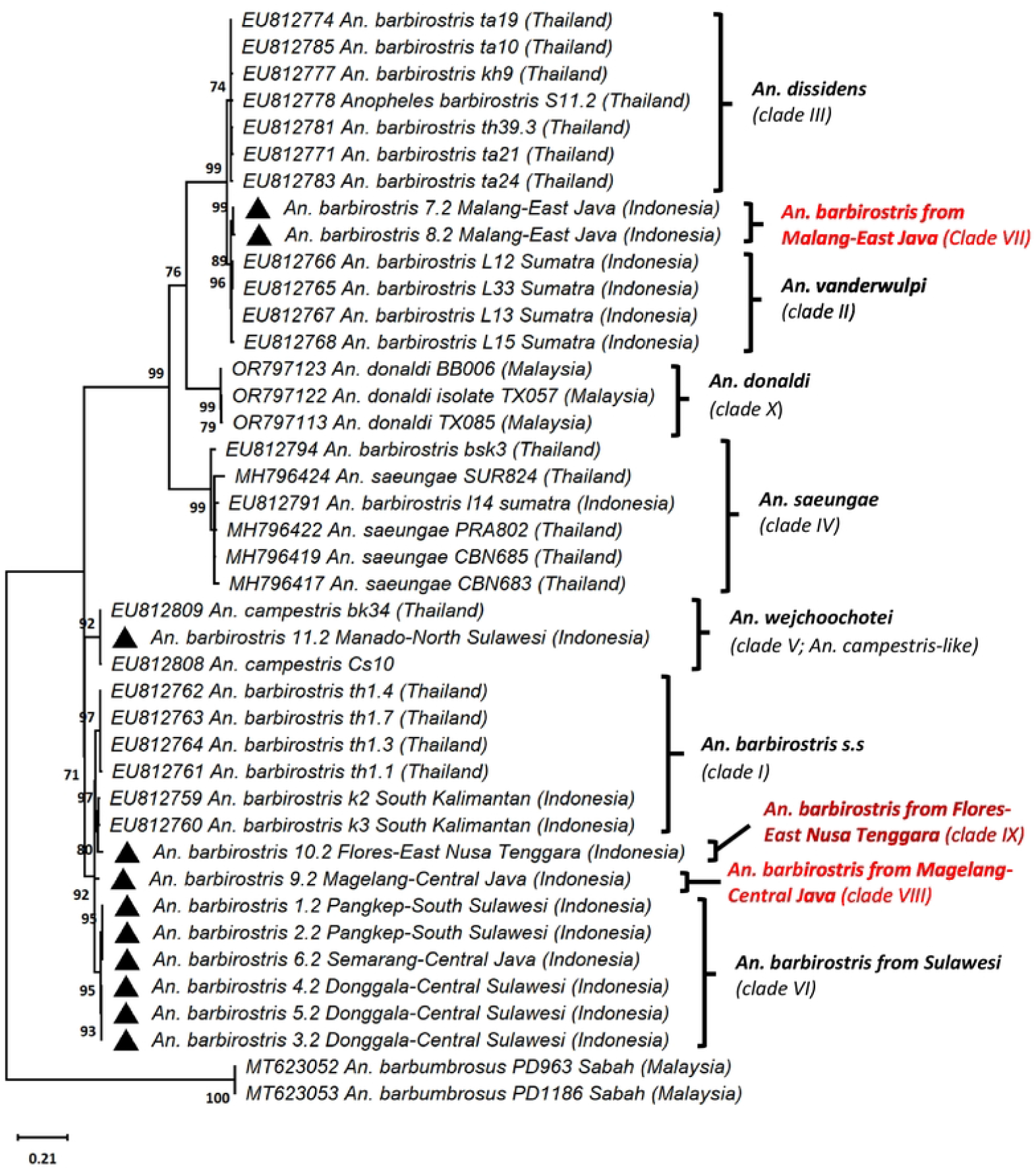
Phylogenetic analysis of the ITS2 sequences of *An. barbirostris* complex. Tree rooted using *Anopheles barbumbrosus* as outgroup. The field collected samples are identified with black triangle. The phylogenetic tree was constructed using the maximum likelihood (ML) method with the Kimura -2 evolutionary model in Mega X. Bootstraps were tested at 1,000 replicates.

### Phylogenetic analysis of the Cytochrome Oxidase I gene (COI)

The analysis of the COI sequences to identify the maternal lineage revealed that all samples and references from the GenBank belonged to three lineages of the *An. barbirostris* complex (Fig. 3). Lineage 1 comprised two GenBank sequences of *An. campestris* from Thailand (EU797276, EU797280), one *An. barbirostris* from Sri Lanka (KC791439), 5 GenBank sequences of *An. barbirostris* from Malang-East Java (JX268791, JX268793, JX268797-JX268798, JX268802), one GenBank sequence of *An. vanderwulpi* from Malang-East Java (JX268804), and also included five different sequences of 106 specimens of *An. barbirostris* from Flores-East Nusa Tenggara (FLORES-S1, FLORES-S3, FLORES-R8, FLORES-R9, 9.2), four from Central Java (10.2, JAVA-J7, JAVA-S8, JAVA-J10), and one from Manado-North Sulawesi (11.2).

**Figure 3.**
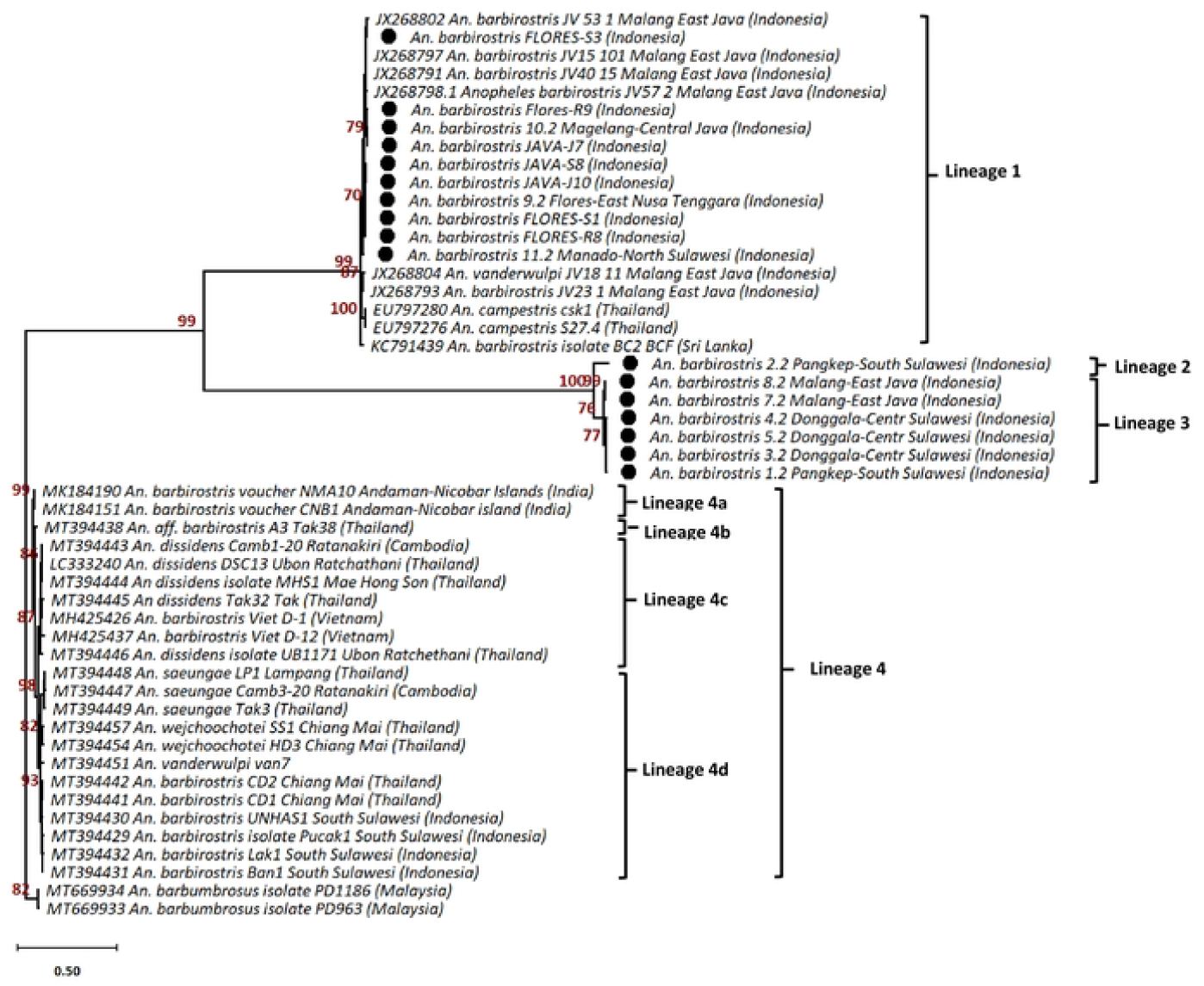
Phylogenetic analysis of the COI sequences. Tree rooted using *Anopheles barbumbrosus* as outgroup. All field collected samples are identified with black dot. Phylogenetic trees were constructed with the General Time Reversible (GTR)+1 model was constructed in Mega X. Bootstraps were tested with 1,000 replicates to assess the reliability of the trees.

Lineage 2 consists of only one specimen of *An. barbirostris* from Pangkep-South Sulawesi (sample 2.2). While six specimens of *An. barbirostris* from Malang-East Java (samples 7.2, 8.2), Donggala-Central Sulawesi (samples 3.2, 4.2, 5.2), and Pangkep-South Sulawesi (sample 1.2) formed lineage 3 (ML bootstrap value 100%).

A branch point pattern indicating lineage 4 is separated into four sub-lineages. Sub-lineage 4a included two GenBank sequences of *An. barbirostris* from the Andaman-Nicobar Islands in India (MK184151, MK184190). Sub-lineage 4b comprised only *An. aff. barbirostris* from Thailand (MT394438). While sub-lineage 4c consists of five GenBank sequences of *An. dissidens*, one from Cambodia (MT394443) and four from Thailand (MT394444, MT394445, MT394446, LC333240), as well as two GenBank sequences of *An. barbirostris* from Vietnam (MH425426, MH425437). Sub-lineages 4d includes GeneBank sequences of *An. barbirostris* from Chiang Mai-Thailand (MT394441-MT394442) and South Sulawesi-Indonesia (MT394429-MT394432), *An. wejchoochotei* from Chiang Mai-Thailand (MT394454, MT394457), *An. saeungae* from Cambodia (MT394447) and Thailand (MT394448-MT394449), and *An. vanderwulpi* var.7 (MT394451) from Java, Indonesia. Two sequences of *An. barbumbrosus* from GenBank (MT669933-MT669934), originated from Malaysia, were used as the outgroup for the rooted tree (Fig. 3).

Indonesian specimens of *An. barbirostris* within the lineage 1 cluster displayed some genetic variability of their COI sequences. *An. barbirostris* from Malang-East Java (JX268791, JX268793, JX268797-JX268798, JX268802) presented a percentage of divergence ranging between 0.2%-0.7%, while genetic distances ranging between 0.6-0.8% were observed with *An. vanderwulpi* from Malang-East Java (JX268804). The COI sequence divergence of the Indonesian samples with *An. campestris* from Thailand (EU797276, EU797280) and *An. barbirostris* from Sri Lanka (KC791439) (Additional file 2) was 0.5% and 0.6-0.7%, respectively.

## Discussion

*Anopheles barbirostris* s.l. has been previously reported in Indonesia as a complex comprising four distinct populations to which species names have now been given, i.e. *An. barbirostris* s.s, *An. saeungae, An. vanderwulpi*, and *An. barbirostris* from Sulawesi [3,17,18]. However, this simply corresponds to the definition of a metapopulation or interfertile populations, in other words four species sharing most of their genetic background but also displaying significantly different genotypic and phenotypic traits. In this work, we report that a total of nine distinct populations can be identified in Indonesia within this *An. barbirostris* metapopulation or complex as commonly referred to in the *Anopheles* classification [2]. *Anopheles wejchoochotei* is a new record within the Indonesian *An. barbirostris* complex. Previously, this latter population from Manado in North Sulawesi was identified as *An. campestris*, although it is not considered a malaria and lymphatic filariasis vector [9,20,31,32]. Thus, while *An. wejchoochotei* had only been reported in Thailand [7,8,13], this study shows that its geographic distribution extends to Indonesia. Furthermore, two specimens of *An. barbirostris* collected from Malang City in East Java (samples 7.2, 8.2) have also been suspected to represent a separate population. They display their closest relationship to *An. vanderwulpi* compared to other members of the *An. barbirostris* complex. This study also revealed that four specimens of *An. barbirostris* from Sulawesi (Pangkep-South Sulawesi - samples 1.2, 2.2, Donggala-Central Sulawesi – samples 4.2, 5.2), and Semarang-Central Java (sample 6.2) formed a distinct clade VI from other *An. barbirostris* specimens from Thailand and Indonesia. They represent thus another distinct population within the *An. barbirostris* complex. As a consequence, previous reports identified *An. barbirostris* from Sulawesi as a distinct species [9,16,18]. Other differing traits linked to the metapopulation structure correspond to variations in bionomics such as behavioral and biological traits as well as vectorial competence. These traits are essential to investigate for implementing efficient vector control strategies. In Sulawesi, *An. barbirostris* plays an important role as a malaria and lymphatic filariasis vector [9,22]. Garjito et al. in 2004 reported that *An. barbirostris* in Donggala, Central Sulawesi, typically feeds outdoors during the night, and female mosquitoes predominantly bite humans from early dusk until midnight [33]. Even though Lien et al. in 1977 reported an anthropophilic form of *An. barbirostris* in Sulawesi, further investigations are needed to describe the various biting behaviors in areas where distinct populations are found [34]. However, these analyses have not been addressed in this work which was devoted to deciphering the population structure as a primary but essential step. The characterization of the bionomics of each population will be the next step.

In this work, a distinct population of *An. barbirostris* has also been identified from Magelang in Central Java, located in the Menoreh hills, a place where a novel species of *An. maculatus* was previously found [35,36]. The existence of distinct populations of *An. barbirostris* in Magelang is inseparable from changes in the continental shelf (2.58 Myrs to 11,700 years ago) that occurred during the Pleistocene Epoch, which resulted in increased volcanism in Central and Eastern Java. Menoreh hills have been previously reported as a separated and preserved area from surrounding volcanic destruction, which served as a relic forest refuge [35,37]. Although this population was not confirmed as a malaria vector in the Java Island, Widiarti et al. reported that *An. barbirostris* specimens from Purworejo, Menoreh Hills, were positive for *P. vivax* infections both by CSP-ELISA and PCR methods [38].

*Anopheles barbirostris* has been documented as a major vector for malaria and lymphatic filariasis in East Nusa Tenggara [21]. Both *P. falciparum* and *P. vivax* were detected in Flores and Adonara Island in the eastern Lesser Sundas [19,21]. This taxon has also been incriminated as vector of lymphatic filariasis in eastern Lesser Sundas (*Brugia timori*) [39]. The existence of eight distinct Barbirostris populations, recorded in this study, provides valuable information on the diversity of malaria and lymphatic filariasis vector species in Indonesia, particularly in East Nusa Tenggara.

Currently, Indonesia is on the way to eliminating lymphatic filariasis and malaria, which are targeted to be achieved by 2030. However, several regions still face problems related to the high transmission of lymphatic filariasis and malaria, especially in Sulawesi and East Nusa Tenggara where *An. barbirostris* acts as the main vector [32,40,41]. Until 2019, the progress rate of lymphatic filariasis elimination in the provinces of Central Sulawesi, Southeast Sulawesi, West Sulawesi, Gorontalo, and South Sulawesi was significant and reached 44% (4/9), 50% (6/12), 50% (2/4), 67% (4/6), and 75% (3/4) of all districts/municipalities, respectively. However, in the province of East Nusa Tenggara, the lymphatic filariasis elimination program has only reached 17% (18/3) of the total districts/municipalities in the area. On the contrary, North Sulawesi is the only province in Sulawesi where all districts/cities are non-endemic for lymphatic filariasis [32].

Moreover, the Ministry of Health of Indonesia also noted that the progress of malaria elimination in the provinces of Gorontalo, Central Sulawesi, North Sulawesi, Southeast Sulawesi, South Sulawesi, and West Sulawesi had reached 66.7% (4/6), 61.5% (8/13), 60% (9/15), 47.1% (9/17), 16.7% (4/24) and 16.7% (1/6) of all districts/municipalities, respectively. However, a total of 34 districts/municipalities in Sulawesi are still malaria endemic areas, of which 33 districts/municipalities have low endemicity, while one district (located in Southeast Sulawesi) is still a medium endemic area [41]. In 2023, Papua was the province where malaria is most endemic accounting for at least 39.17% of all confirmed malaria cases in Indonesia with an annual total of 163,962 cases. Beside Papua, Central Papua is also one of the highest priority regions or provinces (?) for malaria control. Data from the Ministry of Health of Indonesia revealed that as many as 150,225 cases (35.89%) out of 418,546 malaria cases countrywide in 2023 were reported in this province [42]. The third highest malaria endemic area in Indonesia is West Papua. A total of 43,862 malaria cases (10.47%) were reported in 2023 in this province. East Nusa Tenggara is the highest malaria-endemic area outside Papua island, with a total of 6.968 cases (1.66%) identified in the same year [42]. *Anopheles barbirostris* s.l. has been documented in East Nusa Tenggara, but has never been recorded in Papua and West Papua [9].

Over the years, *An. barbirostris* s.l. became one of the main targets for vector control of lymphatic filariasis and malaria in the Lesser Sunda region (mostly East Nusa Tenggara) and Sulawesi [1,9,21,26,34,40,43]. However, most studies on bionomics and vectorial capacity of *An. barbirostris* s.l. in these areas were done, prior to the implementation of molecular analysis techniques, and based on unreliable morphological identification [5,8]. As a consequence, the diversity of populations and species within the *An. barbirostris* complex was not recognized and vector control efforts were then equalized in a variety of habitats and endemic areas. In contrast, the diversity of bionomics of species within this complex has a real impact on malaria epidemiology, transmission, and control, including vectorial capacity, insecticide resistance, feeding preferences, and resting preferences. This is why vector control efforts were not optimally targeted and the disease transmission is still going on [8,43].

This work offers the opportunity to base bionomics and vector competence studies on well-characterized *An. barbirostris* populations.

## Conclusions

Prior to this study, *An. barbirostris* s.l. in Indonesia was considered as having four distinct species, i.e. *An. barbirostris* s.s. in Java and Kalimantan, *An. vanderwulpi* in Java and Sumatra, *An. saeungae* in Sumatra, and an unknown form of *An. barbirostris* in Sulawesi. This study revealed that *An. barbirostris* s.l. in Indonesia is a more complex metapopulation consisting of at least eight populations. The correct characterization of the target populations is essential to better understand the geographic distribution and respective role in disease transmission of each member of the *An. barbirostris* complex. This is particularly true in areas where the transmission is still active, as well as in low transmission areas where the disease is difficult to eliminate.

## Acknowledgments

The authors are grateful for the support from all staff of the Department of Parasitology, Faculty of Medicine, Public Health, and Nursing, Gadjah Mada University, Yogyakarta, Indonesia and all staff and researchers at Human-Animal Interface Laboratory, National Research and Innovation Agency, Salatiga, Indonesia.

## Author contributions

### Conceptualization

Tri Baskoro Tunggul Satoto, Triwibowo Ambar Garjito, Roger Frutos, Sylvie Manguin

### Formal analysis

Tri Baskoro Tunggul Satoto, Triwibowo Ambar Garjito, Roger Frutos, Sylvie Manguin

### Funding acquisition

Tri Baskoro Tunggul Satoto, Triwibowo Ambar Garjito, Sylvie Manguin

### Investigation

Tri Baskoro Tunggul Satoto, Triwibowo Ambar Garjito, Shinta, Soleman Landi

### Methodology

Tri Baskoro Tunggul Satoto, Triwibowo Ambar Garjito, Roger Frutos, Sylvie Manguin

### Project administration

Tri Baskoro Tunggul Satoto, Shinta

### Ressources

Tri Baskoro Tunggul Satoto, Triwibowo Ambar Garjito, Soleman Landi

### Software

Triwibowo Ambar Garjito

### Supervision

Tri Baskoro Tunggul Satoto, Roger Frutos, Sylvie Manguin

### Validation

Triwibowo Ambar Garjito, Roger Frutos, Sylvie Manguin

### Visualisation

Triwibowo Ambar Garjito, Roger Frutos, Sylvie Manguin

### Writing –original draft

Triwibowo Ambar Garjito, Roger Frutos, Sylvie Manguin

### Writing –review & editing

Triwibowo Ambar Garjito, Sylvie Manguin

## Data Availability

All data supporting the conclusions of this article are included within the article and supporting materials.

